# Access to emotional memories: Evidence for a vagal route to boost memory retrieval using non-invasive taVNS

**DOI:** 10.1101/2025.04.17.649268

**Authors:** Manon Giraudier, Carlos Ventura-Bort, Mathias Weymar

## Abstract

Remembering emotionally significant events is critical for the organisms’ behavior and survival. While the mechanisms underlying formation of such memories are well understood and closely linked to the vagus nerve and the brain’s arousal system, less is known about its contribution to memory retrieval. The current study tested whether non-invasive transcutaneous auricular vagus nerve stimulation (taVNS) applied during a long-term recognition memory task influenced retrieval of unpleasant and neutral scenes that have been incidentally encoded one week earlier. Results showed that taVNS, compared to a sham condition, selectively enhanced the recollection of unpleasant images. Despite a modest effect size (*d = 0*.*33*), these findings provide first human evidence for a direct link of the vagus nerve in emotional memory retrieval, thus extending current theoretical models of emotional memory retrieval and opening a new pathway for memory modulation and non-invasive therapeutic interventions.

## Introduction

Remembering emotionally significant events is essential in various aspects of everyday life, as emotional memories guide decision-making, shape social interactions, and influence mental well-being (LaBar & Cabeza, 2006; Dolcos et al., 2020). While the mechanisms underlying the formation of such powerful and enduring memories are well understood, less is known about how these memories are retrieved. Understanding these processes, however, is important not only for basic research but also for clinical applications, for example, in post-traumatic stress disorder (PTSD) and neurodegenerative diseases such as Alzheimer’s disease, where dysregulation of noradrenergic function may lead to memory distortions or deficits (Marien et al., 2004; O’Donnell et al., 2004; Dahl et al., 2019; Holland et al., 2021). In aging populations, a decline in noradrenergic integrity has been associated with reduced emotional memory selectivity, potentially affecting adaptive decision-making (Mammarella et al., 2016; Dahl et al., 2023). Identifying mechanisms that regulate memory retrieval could therefore provide novel strategies to mitigate cognitive deficits and enhance memory function.

One promising target for modulating retrieval processes is the vagus nerve, a central mediator between peripheral physiological states and central arousal mechanisms. Through afferent projections to the nucleus of the solitary tract (McGaugh et al., 2000; Schwabe, 2022), the vagus nerve influences the locus coeruleus-noradrenaline (LC-NA) system, which modulates the amygdala and medial temporal lobe (MTL), structures critical for memory formation (McIntyre et al., 2012). Specifically, the amygdala-hippocampus interaction facilitates the encoding and consolidation of emotional memory, while noradrenaline modulates these processes (Strange & Dolan, 2004; Weymar & Hamm, 2013). While the LC-NA system is well recognized for its role in encoding and consolidation of emotional memories (Mather et al., 2016), its involvement in retrieval is less established. Evidence from animal studies suggests that direct stimulation of the LC during retrieval might enhance memory performance, alleviating forgetting through beta-adrenergic receptor activation, also highlighting the role of noradrenaline in modulating retrieval processes (Devauges & Sara, 1991). In humans, functional neuroimaging data indicate that successful retrieval of neutral stimuli encoded in emotional contexts involves LC activity, with functional interactions between the LC and amygdala playing a key role in this process (Sterpenich et al., 2006). These two findings suggest that the LC-NA arousal system may also be critical during retrieval by influencing attentional and mnemonic processes. Transcutaneous auricular vagus nerve stimulation (taVNS), a non-invasive method that stimulates the auricular branch of the vagus nerve, has emerged as a promising tool for modulating the LC-NA system (Kraus et al., 2013; Yakunina et al., 2017; Sharon et al., 2021) and its associated memory functions. Recent studies provide evidence that taVNS during encoding enhances recollection-based memory (i.e., the retrieval of specific contextual details associated with the highest level of memory confidence) (Giraudier et al., 2020), particularly for emotionally arousing contents (Ventura-Bort et al., 2021; Ventura-Bort et al., 2025), a mnemonic process related to amygdala and hippocampal activity (Dolcos et al., 2005, 2010). Whether vagal stimulation during retrieval, via LC-NA activation, also improves recollection-based memory for emotional contents, however, is not known so far.

The present study aimed to uncover the mechanisms underlying memory retrieval, thereby contributing to current theoretical frameworks in cognitive neuroscience. Specifically, we investigated whether non-invasive taVNS, applied during a long-term memory recognition task, influenced retrieval of emotional and neutral scenes that had been incidentally encoded one week earlier. Based on prior findings we expected a memory advantage for emotionally salient stimuli, with this enhancement being driven by recollection-based processes. If the vagus nerve - via activation of the LC-NA system is involved in memory retrieval (c.f., Devauges & Sara, 1991), we expect that taVNS would enhance recollection-based retrieval, particularly for unpleasant images.

## Materials and Methods

### Sample

A total of 80 healthy students from the University of Potsdam participated in the study in exchange of course credits^1^. All participants had normal or corrected-to-normal vision and were German speakers (at least C1 levels). Exclusion criteria included neurological or psychiatric disorders, brain surgery, undergoing medication or drug use, pregnancy, a history of migraine and/or epilepsy, cardiac diseases, metal pieces in the body (i.e., a pacemaker), and active implants or physical alterations in the ear (e.g., a cochlear implant). Based on these criteria, five participants were excluded due to reported diagnoses, including anxiety disorders, depression, attention-deficit/hyperactivity disorder and rat phobia. The final sample consisted 75 participants, who were randomly assigned to either the taVNS group (*N = 38, 33 female, 5 male, M*_*age*_ *= 21*.*1, SD*_*age*_ *= 2*.*71*) or the sham stimulation group (*N = 37, 31 female, 6 male, M*_*age*_ *= 21*.*8, SD*_*age*_ *= 3*.*79*). Each individual provided written informed consent for the study protocol and the application of taVNS in the context of an emotional picture recognition memory paradigm was approved by the Ethics Committee of the University of Potsdam. All data are publicly available (https://osf.io/rqs6p/).

### Experimental procedure

A 2-day randomized, single-blinded, between-subject design was employed, with participants randomly assigned to either receive taVNS or sham stimulation. The experiment was conducted over two separate sessions (seven days apart): an encoding session and a recognition session (see Figure 1). Stimulation was only administered during the recognition session.

**Figure 1:**
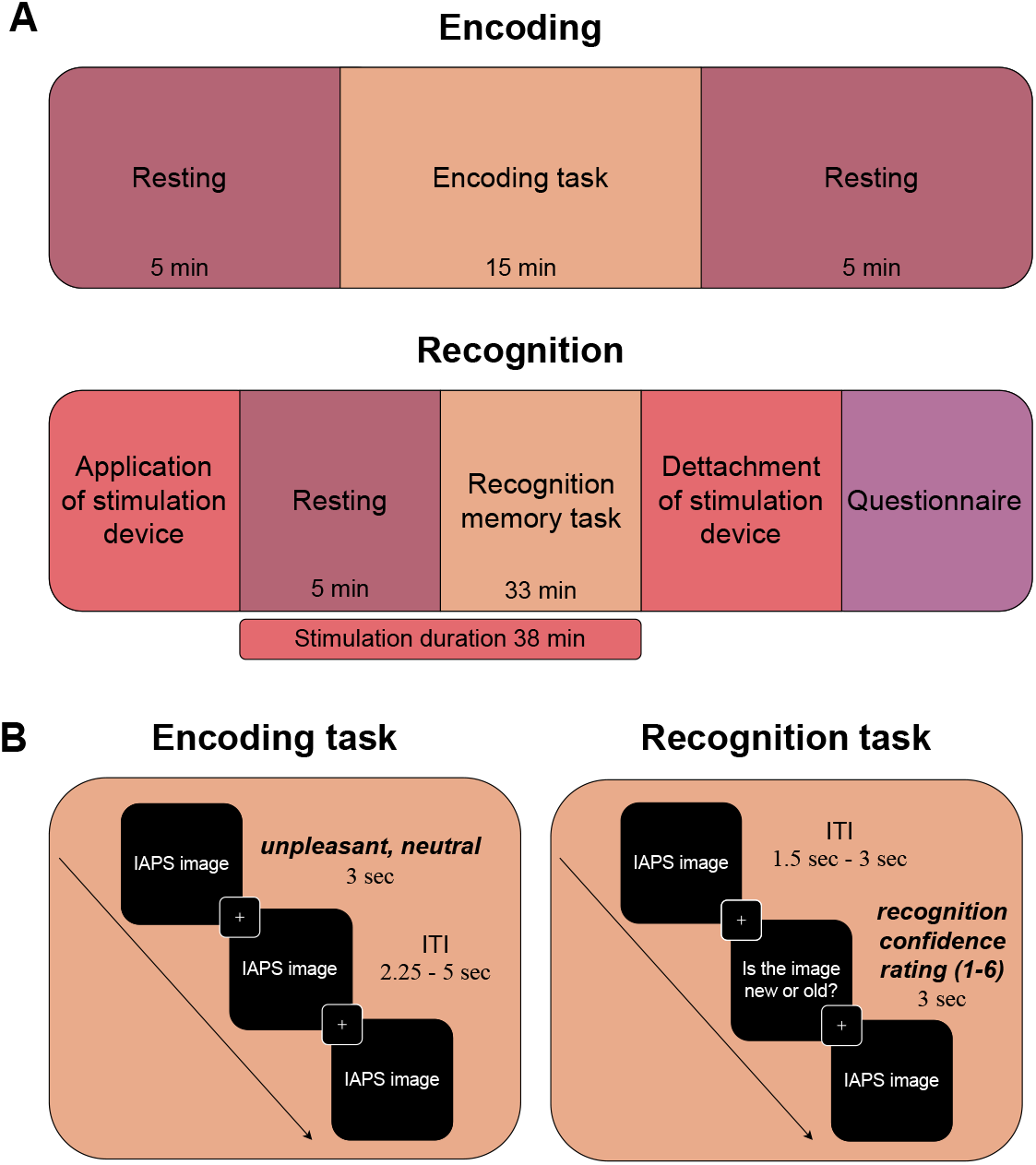
Experimental design consisting of two sessions conducted seven days apart. **(A)** The first session included an encoding task, with 5 min resting phases before and after the task. The second session consisted of a recognition memory task, preceded by a 5 min resting phase with either taVNS or sham stimulation and followed by a side effects questionnaire. **(B)** During encoding, participants attentively viewed neutral and unpleasant images. In the recognition task, participants were presented with previously seen images and new images and rated their memory confidence on a 6-point scale, ranging from 1 (*definitely new*) to 6 (*definitely old*).

During the encoding session, participants were seated in front of a computer screen and presented with a total of 120 images, consisting of 60 neutral and 60 unpleasant images. Each image was displayed for 3000 ms, with an inter-trial interval between 2500 and 5000 ms. Picture presentation was pseudorandomized with no more than two images from the same category presented consecutively. Before and after the task, participants underwent 5 min resting phases without any image presentation. During the task, participants were instructed to view the images attentively, without being informed about the subsequent memory test, ensuring incidental encoding. One week after the encoding session, participants returned to the laboratory for the recognition session. Upon arrival, the stimulation electrodes were attached to the left ear of participants, and the stimulation intensity was calibrated according to individual sensitivity (for detailed protocol see **Transcutaneous Auricular Vagus Nerve Stimulation** section). Following calibration, participants received 5 min of stimulation without any task. Subsequently, participants performed a recognition memory task, which lasted 33 minutes and included all of the 120 previously encoded images randomly intermixed with 120 new images, resulting in a total of 240 trials. Each image was displayed on the screen for 3000 ms, preceded by a fixation cross. After each image, participants were asked to assess their recognition memory allowing to differentiate between familiarity and recollection using a 6-point confidence scale, with 1 indicating the picture is *definitely new*, 2 indicating the picture is *probably new*, 3 indicating the picture is *perhaps new*, 4 indicating the picture is *perhaps old*, 5 indicating the picture is *probably old* and 6 indicating the picture is *definitely old*.

After the completion of the task, the stimulation electrodes were removed, and participants completed a side-effects questionnaire (adapted from Jacobs et al., 2015), for which they had to indicate on a 7point Likert scale (1 being *not at all* and 7 being *very much*), how much they had experienced headaches, nausea, dizziness, neck pain, muscle contractions in the neck or face, stinging sensations under the electrode, skin irritation at the ear, concentration fluctuations, mood changes, and unpleasant feelings.

### Stimulus material

A total of 240 images selected from the *International Affective Picture System* (IAPS) (Lang et al., 2008), the *Nencki Affective Picture System* (NAPS) (Marchewka et al., 2014), and the *EmoMadrid Set* (Carretié et al., 2019) were used in the experiment. The images were preselected based on their standardized valence and arousal ratings, and included 120 neutral contents (e.g., buildings, neutral landscapes, human faces with neutral expressions; *M*_*valence*_ *= 5*.*23, SD*_*valence*_ *= 0*.*35; M*_*arousal*_ *= 4*.*40, SD*_*arousal*_ *= 0*.*84*) and 120 unpleasant contents (e.g., mutilations, attacks, accidents; *M*_*valence*_ *= 2*.*30, SD*_*valence*_ *= 0*.*71; M*_*arousal*_ *= 7*.*29, SD*_*arousal*_ *= 0*.*98*). Images were counterbalanced across participants by creating eight different image lists. Each list was divided into four sets, with each set containing 30 neutral and 30 unpleasant images. These sets were matched in terms of valence, arousal, and content. During the retrieval task, each set was used either as a previously seen or a novel image, and the assignment of sets to experimental conditions was counterbalanced across participants. Each participant was randomly assigned to one of the eight lists, preventing order or list effects.

### Transcutaneous auricular vagus nerve stimulation

The stimulation device consisted of two titanium electrodes mounted on a holder resembling in-ear headphones, connected to a battery-operated stimulation unit (CMO2, Cerbomed, Erlangen, Germany). In the taVNS condition, the electrodes were positioned in the left cymba conchae, an area innervated exclusively by the auricular branch of the vagus nerve (ABVN) (Peuker & Filler, 2002). As in previous studies using taVNS (e.g., Giraudier et al., 2020; Ventura-Bort et al., 2021), for sham condition, the electrodes were placed on the center of the left earlobe, an area known to be free of vagal innervation (Peuker & Filler, 2002). Electrical stimulation was applied in alternating 30-second cycles of stimulation and rest. The device delivered stimulation with a pulse width of 250 µs and a frequency of 25 Hz. Stimulation intensity was individually calibrated to be above the sensory threshold but below the pain threshold (cf., Ventura-Bort et al., 2021). The stimulation intensity differed between the taVNS (*M = 1*.*12, SD = 0*.*615*) and the sham conditions (*M = 1*.*49, SD = 0*.*481*), *t(68*.*10) = 2*.*93, p = 0*.*005*. However, this difference was not expected to influence the results, as stimulation intensity has generally shown no significant effects on physiological or behavioral outcomes (e.g., Borges et al., 2019; Keatch et al., 2023; Janner et al., 2018). As expected, the inclusion of stimulation intensity as a covariate in the statistical models also did not change the reported memory effects (see Appendix A).

### Statistics

All statistical analyses were carried out in the R environment (version 4.3.0.).

#### Self-report measures

To test whether the reported side effects differed between taVNS and sham stimulation, t-tests were performed for each reported subjective symptom, separately.

#### Recognition memory performance

To assess the effects of taVNS on recognition memory performance, the discrimination index Pr was defined as *Pr* = *p*(*hit*) − *p*(*false alarm*), where higher Pr values indicate better discrimination between old and new images. The response bias index Br was calculated as 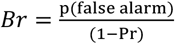, where Br > 0.5 reflects a liberal response criterion (i.e., a bias toward responding *old*), and lower values indicate a conservative response bias (Snodgrass & Corwin, 1988). Responses were categorized as *old* if rated 4-6 on the confidence scale and *new* if rated 1-3. The Pr and the Br index were analyzed using a linear mixed-effects model, with fixed effects of *Stimulation* (taVNS vs. sham) and *Category* (unpleasant vs. neutral), including their interaction and a random intercept for participants.

#### Recognition memory performance based on confidence ratings

Memory performance was divided according to responses on the 6-point confidence scale to differentiate between recollectionand familiarity-based retrieval processes. Recollection-based memory was identified using responses with a rating of 6 (*definitely old*), while familiarity-based memory was calculated using responses with ratings of 4 (*perhaps old*) and 5 (*probably old*). The Pr index was analyzed using a linear mixed-effects model, with fixed effects of *Stimulation* (taVNS vs. sham), *Category* (unpleasant vs. neutral), and *Memory* (familiarity vs. recollection), including their interactions. Participant intercepts and slopes for *Memory* were modeled as random effects to account for individual variability in response patterns. The selected random-effect structure included theoretically relevant variance components and was supported by the data (c.f., Bates, 2015). Data points falling outside 1.5 times the interquartile ranges above the third quartile or below the first quartile (1% of the data) were excluded from further analysis (Wickham & Stryjewski, 2011).

#### Recollection index

To further investigate the role of recollection-based memory for unpleasant and neutral images and its modulation by Stimulation, a recollection index (RI) was calculated. This index represented the advantage of recollection-based memory over familiarity-based memory for each Category, *RI*_*Unpleasant*_ = *Pr*_*Recollection*_*Unpleasant*_ − *Pr*_*Familiarity*_*Unpleasant*_ and, *RI*_*Neutral*_ = *Pr*_*Recollection*_*Unpleasant*_ − *Pr*_*Familiarity*_*Unpleasant*_. By computing this difference, we isolated the extent to which recollection processes contributed to discrimination performance relative to familiarity-based memory, providing a clear measure of the recollection advantage for both neutral and unpleasant images. The RI was analyzed using a linear mixed-effects model, with fixed effects of *Stimulation* (taVNS vs. sham) and *Category* (unpleasant vs. neutral), including their interaction and a random intercept for participants.

## Results

### Side effects of stimulation

As expected, participant’s subjective ratings revealed minimal side effects associated with the stimulation (*M = 1*.*97, SD = 1*.*26*) and there was no indication of significant differences between taVNS and sham stimulation for any of the assessed side effects (*ps > 0*.*19*; see Appendix B) (c.f., Giraudier et al., 2025).

### Recognition memory performance

The behavioral results for the recognition memory task, as a function of *Category* and *Stimulation*, are summarized in Table 1. As expected, a significant main effect of *Category, β = 0*.*10, SE = 0*.*023, t(73) = 4*.*28, p* < *0*.*001*, indicated significantly higher Pr values for unpleasant images compared to neutral images. This effect, however, was not significantly influenced by *Stimulation, β = 0*.*01, SE = 0*.*03, t(73) = 0*.*40, p = 0*.*69*, indicating that taVNS did not further enhanced the memory advantage for unpleasant images. The main effect of *Stimulation, β = 0*.*02, SE = 0*.*043, t(97*.*47) = 0*.*38, p = 0*.*71*, was also not significant.

**Table 1.** Mean (standard deviation) of behavioral indices for unpleasant and neutral images encoded under taVNS and sham stimulation.

|  | taVNS |  | Sham |  |
| --- | --- | --- | --- | --- |
|  | Unpleasant | Neutral | Unpleasant | Neutral |
| **Outcome rates** |  |  |  |  |
| Hits | 0.82 (0.12) | 0.64 (0.17) | 0.79 (0.12) | 0.63 (0.18) |
| False alarms | 0.24 (0.12) | 0.17 (0.12) | 0.24 (0.16) | 0.18 (0.12) |
| **Discrimination index** |  |  |  |  |
| Pr | 0.58 (0.16) | 0.47 (0.19) | 0.55 (0.18) | 0.45 (0.21) |
| **Response bias index** |  |  |  |  |
| Br | 0.59 (0.24) | 0.33 (0.20) | 0.51 (0.22) | 0.34 (0.18) |
| **Confidence rates** |  |  |  |  |
| Familiarity-based Pr | 0.24 (0.14) | 0.29 (0.10) | 0.26 (0.11) | 0.29 (0.12) |
| Recollection-based Pr | 0.58 (0.20) | 0.35 (0.20) | 0.53 (0.17) | 0.35 (0.17) |
| **Recollection index** |  |  |  |  |
| RI | 0.51 (0.29) | 0.22 (0.26) | 0.42 (0.22) | 0.21 (0.21) |

Moreover, for Br, participants demonstrated a more liberal response bias (i.e., higher Br values) for unpleasant images compared to neutral images, *β = 0*.*17, SE = 0*.*03, t(73) = 5*.*90, p* < *0*.*001*. Although the main effect of *Stimulation* was not significant, *β =* −*0*.*01, SE = 0*.*05, t(104*.*38) = 0*.*17, p = 0*.*87, the* interaction between *Stimulation* and *Category* approached significance, *β = 0*.*08, SE = 0*.*04, t(73) = 1*.*94, p = 0*.*056*, suggesting a trend toward a more liberal response bias for unpleasant images under taVNS compared to sham stimulation.

### Recognition memory based on recollection and familiarity

Unpleasant images were generally associated with significantly lower Pr values compared to neutral images, *β =* −*0*.*05, p = 0*.*011*. Additionally, a strong main effect of *Memory* was observed, *β = 0*.*21, p* < *0*.*001*, indicating that recollection-based memory was associated with generally better discrimination compared to familiarity-based memory. The main effect of *Stimulation* was not significant, *β = 0*.*01, p = 0*.*861*.

A significant interaction between *Category* and *Memory*, however, showed that the advantage of recollection over familiarity was greater for unpleasant images than for neutral images, *β = 0*.*20, p* < *0*.*001*, replicating prior findings. Post hoc testing further supported this effect for both sham stimulation, *β =* −*0*.*15, p* < *0*.*001*, and taVNS, *β =* −*0*.*21, p* < *0*.*001*, with a stronger enhancement under taVNS. Familiarity-based memory showed a smaller, reverse pattern, with neutral images outperforming unpleasant images, particularly under taVNS, *β = 0*.*09, p* < *0*.*001*. However, no interaction was found between *Category* and *Stimulation, β =* −*0*.*04, p = 0*.*19*, nor between *Memory* and *Stimulation, β = 0*.*00, p = 0*.*996*.

Critically, a significant three-way interaction between *Stimulation, Memory*, and *Category, β = 0*.*09, p = 0*.*019*, suggested that taVNS modulated memory performance specifically for unpleasant images during recollection compared to sham (see Figure 2). Post-hoc testing revealed no significant difference for unpleasant images between taVNS and sham for recollection-based memory, *β =* −*0*.*06, p = 0*.*14*. The effect size for the difference between stimulation conditions was small, *d = 0*.*33*, consistent with a modest effect of taVNS on recollection for unpleasant images (Funder & Ozer, 2019). Across both stimulation conditions, recollection-based memory was significantly better than familiarity-based memory for all image categories. The effect was stronger for unpleasant images under taVNS, *Unpleasant*_*Sham*_: *β =* −*0*.*41, p* < *0*.*001, Unpleasant*_*taVNS*_: *β =* −*0*.*51, p* < *0*.*001*, aligning with the hypothesis that recollection processes play a greater role in memory for unpleasant images, particularly under taVNS. Moreover, significant variability across participants was observed for the intercept (*SD = 0*.*09*) and the slope for *Memory* (*SD = 0*.*22*), with a negative correlation between them (*r =* −*0*.*75*).

**Figure 2:**
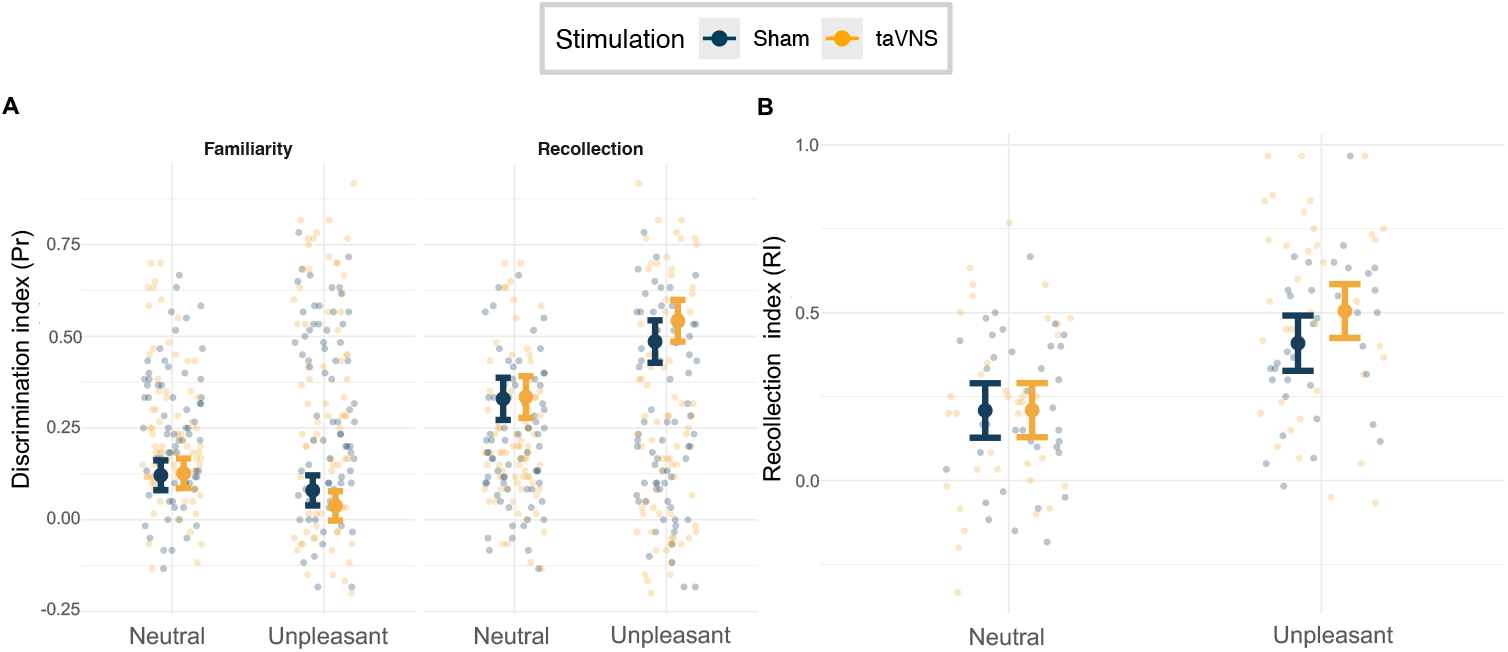
**A.** The y-axis depicts the discrimination index (Pr), while the x-axis represents the *Category* (unpleasant and neutral images), analyzed for familiarity-based and recollection-based memory under taVNS (orange) and sham stimulation (blue). **B.** The y-axis shows the recollection index (RI), with the x-axis indicating *Category* (unpleasant and neutral images), comparing taVNS (orange) and sham stimulation (blue).

### Recollection index

The analysis of the RI showed that unpleasant images generally exhibited a greater recollection advantage compared to neutral images, *β = 0*.*20*, SE = 0.03, t(71.59) = 6.24, *p* < *0*.*001*. Although *Stimulation* alone did not significantly influence the recollection advantage, *β = 0*.*001, SE = 0*.*06, t(98*.*82) = 0*.*02, p = 0*.*987*, a significant interaction between *Stimulation* and Category, *β = 0*.*09, SE = 0*.*05, t(71*.*84) = 2*.*02, p = 0*.*047*, indicated that this advantage for unpleasant images was particularly enhanced under taVNS, suggesting that taVNS selectively modulated recollection-based memory processes for emotionally arousing (unpleasant) images (see Figure 2).

## Discussion

In this study, we investigated the role of the vagus nerve in memory retrieval providing first human evidence that vagal activation through non-invasive taVNS selectively enhanced the recollection-based retrieval of emotionally salient memories.

### Enhanced recollection memory for emotional contents under taVNS

As expected, we replicated the well-established memory advantage for emotionally salient stimuli, which were better remembered than neutral ones, particularly through recollection-based memory (Bradley et al., 1992; Yonelinas, 2002; Dolcos et al., 2005; Weymar & Hamm, 2013), an effect driven by arousal-mediated LC-NA system activation during memory formation (Sterpenich et al., 2006; Mather et al., 2016; Roozendaal & McGaugh, 2011; Dolcos et al., 2020). Critically, such noradrenergic activation seems to be also important for memory retrieval, as taVNS applied during memory reactivation, seemed to boost memory, particularly for emotionally salient stimuli. This finding supports the hypothesis that taVNS modulates LC-NA system activity via afferent projections to the hippocampus-mediated memory system that supports recollection-based memories (Devauges & Sara, 1991; Sterpenich et al., 2006; Weymar & Hamm, 2013). An important mechanism underlying the vagally-mediated retrieval effects could be the reinstatement of an arousal state during retrieval that mirrors the one experienced during encoding (Yaffe et al., 2014). The LC-NA system plays a crucial role in regulating arousal and attention (Aston-Jones & Cohen, 2005; Sara, 2009), and its activation during retrieval may facilitate access to previously encoded information by re-establishing a neural state similar to that present at encoding (Wing et al., 2015). Moreover, beyond reinstating encoding-related arousal states, taVNS might also enhance memory retrieval by modulating the tonic levels of LC-NA activity. Increased tonic activation of the LC-NA system could improve the signalto-noise ratio in neural processing, thereby optimizing the general efficiency of cognitive operations (c.f. Aston-Jones & Cohen, 2005). Under this framework, the enhanced recollection of unpleasant images under taVNS might be one instance of a broader mechanism wherein LC-NA activity improves task-relevant neural processing. While in this case the goal of the task is to retrieve past experiences, in other contexts, such as attentional control, similar LC-NA modulation may facilitate performance by biasing processing toward relevant stimuli and suppressing irrelevant information (Mather et al., 2016; Ehlers, 2017; Bari et al., 2020). Through its connection to areas critical for memory formation and emotional processing, such as the amygdala and hippocampus, the LC-NA system may further enhance neural processing within these cognitive and affective domains (Sara, 2009; McIntyre et al., 2012), leading to improved recollection memory for emotional content under taVNS. Consistent with this framework, animal findings have shown that LC stimulation facilitates memory retrieval via noradrenergic receptor activation (Devauges & Sara, 1991), supporting the idea that taVNS may exert its effects through similar pathways. While previous studies primarily examined encoding effects of taVNS and reported enhanced emotional recollection memory (Ventura-Bort et al., 2021; Ventura-Bort et al., 2025), the current results highlight its role in retrieval, emphasizing the broader functional significance of LC-NA modulation in emotional memory retrieval.

### Future Directions, limitations

This study provides not only first human evidence that vagal activation via taVNS enhances emotional memory retrieval (which expands prior work on memory formation, e.g., Ventura-Bort et al., 2021), but also points towards a new pathway for memory modulation and non-invasive therapeutic interventions. By demonstrating that taVNS selectively improves recollection memory, our findings hold important implications, for instance, for forensic and clinical applications, such as enhancing the accuracy of eyewitness testimony and facilitating memory performance in neurodegenerative conditions such as Alzheimer’s disease. To further extend the current findings, because we used a recognition memory task, future research should also explore whether taVNS exerts a similar memory improving effect when applied during other tasks, testing explicit (e.g., free recall) or implicit forms of memories. Moreover, a systematic investigation of stimulation timing whether applied during encoding, retrieval, or both - would further clarify its differential effects on memory processes. Disentangling the specific contributions of LC-NA activity to encoding versus retrieval and addressing the absence of direct physiological LC-NA markers would provide a foundation for targeted interventions.

Although our statistical models suggest that the observed effects are robust, future studies should minimize potential between-group variability to ensure that the observed effects are attributable to the stimulation itself rather than pre-existing differences between groups. Additionally, the observed effect sizes were small-to-moderate, indicating a modest but reliable effect of taVNS on recollection for unpleasant images. This highlights the importance of replication studies to further validate and refine our understanding of this effect.

## Conclusion

In conclusion, this study is the first to apply taVNS during memory retrieval and demonstrated that taVNS selectively enhanced recollection memory for emotionally salient (unpleasant) stimuli, supporting its hypothesized role in modulating LC-NA system activity. These findings advance our understanding of retrieval processes and underscore taVNS as a promising tool for investigating LC function in memory, with potential applications for non-invasive therapeutic interventions. More broadly, the results contribute to theoretical models of emotional memory by highlighting the role of arousal-related neuromodulation in shaping retrieval processes.

## Data availability statement

All data and analysis scripts are publicly available (https://osf.io/rqs6p/).

## Footnotes

1 *A posteriori* power analysis was conducted using G*Power (version 3.1.9.7.), based on the resulting effect size (*d* = *0*.*33*, *f* = *0*.*15*, *α* = *0*.*05*) and estimated a power of 0.8 with 80 participants.

## Notes

https://osf.io/rqs6p/

### Competing Interest Statement

The authors have declared no competing interest.

