## Appendix for "Access to emotional memories: Evidence for a vagal route to boost memory retrieval using non-invasive taVNS"

| | | | **Fixed effects** | | |
|---|---|---|---|---|---|
| | **Est** | **SE** | **DF** | **t-value** | **p-value** |
| Intercept | 0.11 | 0.03 | 114.00 | 3.53 | **< 0.001** |
| Stimulation | 0.01 | 0.03 | 116.80 | 0.25 | 0.81 |
| Category | -0.04 | 0.02 | 143.20 | -2.11 | **0.04** |
| Memory | 0.21 | 0.04 | 91.90 | 5.17 | **< 0.001** |
| Stimulation intensity | 0.01 | 0.02 | 72.11 | 0.34 | 0.74 |
| Stimulation:Category | -0.05 | 0.03 | 143.20 | -1.74 | 0.08 |
| Stimulation:Memory | 0.00 | 0.06 | 92.29 | -0.01 | 0.99 |
| Category:Memory | 0.20 | 0.03 | 142.90 | 7.14 | **< 0.001** |
| Stimulation:Category:Memory | 0.10 | 0.04 | 142.70 | 2.57 | **0.01** |
| | | | **Random effects** | | |
| | **Variance** | | **Std. Dev.** | | **Corr.** |
| Participant (Intercept) | 0.01 | | 0.09 | | |
| Memory | 0.05 | | 0.21 | | -0.75 |

**Appendix A:** Results of a linear mixed-effects model (LMM) assessing the effects of *Stimulation*, *Category* and *Memory*, with *Stimulation intensity* included as a covariate. The model was fitted to N = 297 observations from 75 participants. The inclusion of stimulation intensity did not influence the observed memory effects. Confirming the robustness of the primary findings (*Est = estimate, SE = standard error, DF = degrees of freedom, Std. Dev. = standard deviation, Corr. = correlation*).

| | **taVNS** | **Sham** | **p-value** |
|---|---|---|---|
| **Concentration** | 2.78 (1.55) | 3.14 (1.18) | 0.38 |
| **Dizziness** | 1.58 (1.18) | 1.35 (0.79) | 0.33 |
| **Fluctuation of Feelings** | 1.81 (1.41) | 1.95 (1.18) | 0.66 |
| **Headache** | 1.81 (1.31) | 1.62 (0.98) | 0.48 |

| | | | |
|---|---|---|---|
| **Nausea** | 1.32 (0.91) | 1.30 (0.85) | 0.89 |
| **Neck Contraction** | 2.03 (1.17) | 1.68 (1.13) | 0.19 |
| **Neck Pain** | 1.41 (0.64) | 1.43 (1.01) | 0.89 |
| **Skin Irritation** | 1.92 (1.61) | 1.89 (1.66) | 0.93 |
| **Stinging Sensation** | 2.97 (1.83) | 2.73 (1.82) | 0.57 |
| **Unpleasant Feelings** | 2.30 (1.51) | 2.38 (1.44) | 0.81 |

**Appendix B:** Mean subjective rating (standard deviation) for taVNS and sham stimulation. Ratings were scored on a seven-point scale with 1 being *not at all* and 7 being very much.
